# Confidence: A Web App for Cross-Platform Differential Gene Expression Analysis, Gene Scoring, and Enrichment Analysis

**DOI:** 10.1101/2025.06.27.661997

**Authors:** Abhishek Shastry, Benjamin P. Ott, Alex Paterson, Matt Simpson, Kimberly J. Dunham-Snary, Charles C.T. Hindmarch

## Abstract

RNA sequencing (RNA-seq) is used to quantify transcript levels through measurement of nucleotide sequences. To evaluate statistically significant changes in gene expression, transcript counts between samples are compared using differential expression analysis methods. However, three of the most pressing challenges in transcriptomics analyses are: 1) analytical packages produce a distinct number of differentially expressed genes with varied *P*-value and fold-change values; 2) the effective use of these analytical packages requires substantial knowledge of programming and bioinformatics; and 3) there are a lack of intuitive methods to select target genes for further investigation in an unbiased manner. To address these challenges, we developed Confidence, a web-based application to perform simultaneous statistical analysis of RNA-seq count data. Confidence incorporates the Confidence Score (CS), ranging from 1 to 4 to aid in gene prioritization, where 1 represents low confidence and 4 represents high confidence. The Confidence web-based application was designed for rapid and intuitive analysis of standard experimental metadata and gene count inputs. Confidence provided a web-based, ‘wide-net’ approach to differential gene expression analysis. Gene scoring allows for unbiased gene selection and identification of novel genes strongly associated with disease and treatment models across multiple species. Additionally, pathway analysis tools have been integrated so that highly confident genes can be placed into biological context in terms of functions and pathways. Confidence provides a new strategy for target prioritization in RNA-seq analysis and the generation of publication-quality figures.

## Background

The draft of the human genome in 2001 introduced a post-genomic era that promised global resolution of the molecular landscape in both preclinical and clinical research environments *(1)*. To capitalize on this new information, new technologies were developed; among them, the microarray, capable of profiling the simultaneous expression of tens of thousands of genes *(2-4)*. The endpoint of a microarray experiment is the identification of differentially expressed genes between two conditions *(5-7)*. The emergence of various RNA sequencing (RNA-seq) technologies substantially improved genome-wide transcriptomic analyses. Rather than relying on the hybridization of mRNA fragments to a library of prescribed probes, as with microarrays *(3, 4, 8)*, RNA-seq turned the sample into a library, facilitating novel transcript discovery *(9-15)*. This high-throughput sequencing method provides several benefits compared to older transcriptomic analysis methods, including the ability to differentiate between isoforms of the same gene, identify coding and non-coding genes, and detect low-abundance transcripts *(9, 16-18)*. The products of RNA-seq are often short, unaligned reads from fragments of the template material. These sequence data are stored in .FASTA or .FASTQ files. To characterize gene expression, reads must be aligned to a previously compiled, annotated, and species-specific reference genome. As the number of defined genomes increased after the assembly of the human genome, several genome alignment tools also emerged to assess the probability that sequenced reads match known genomic sequences *(19-23)*. After genome alignment, gene counts can be approximated using predictive models assessing the probability of alignment to exon sequences *(24-28)*. A schematic of the Confidence workflow is found in **Figure 1**.

**Figure 1.**
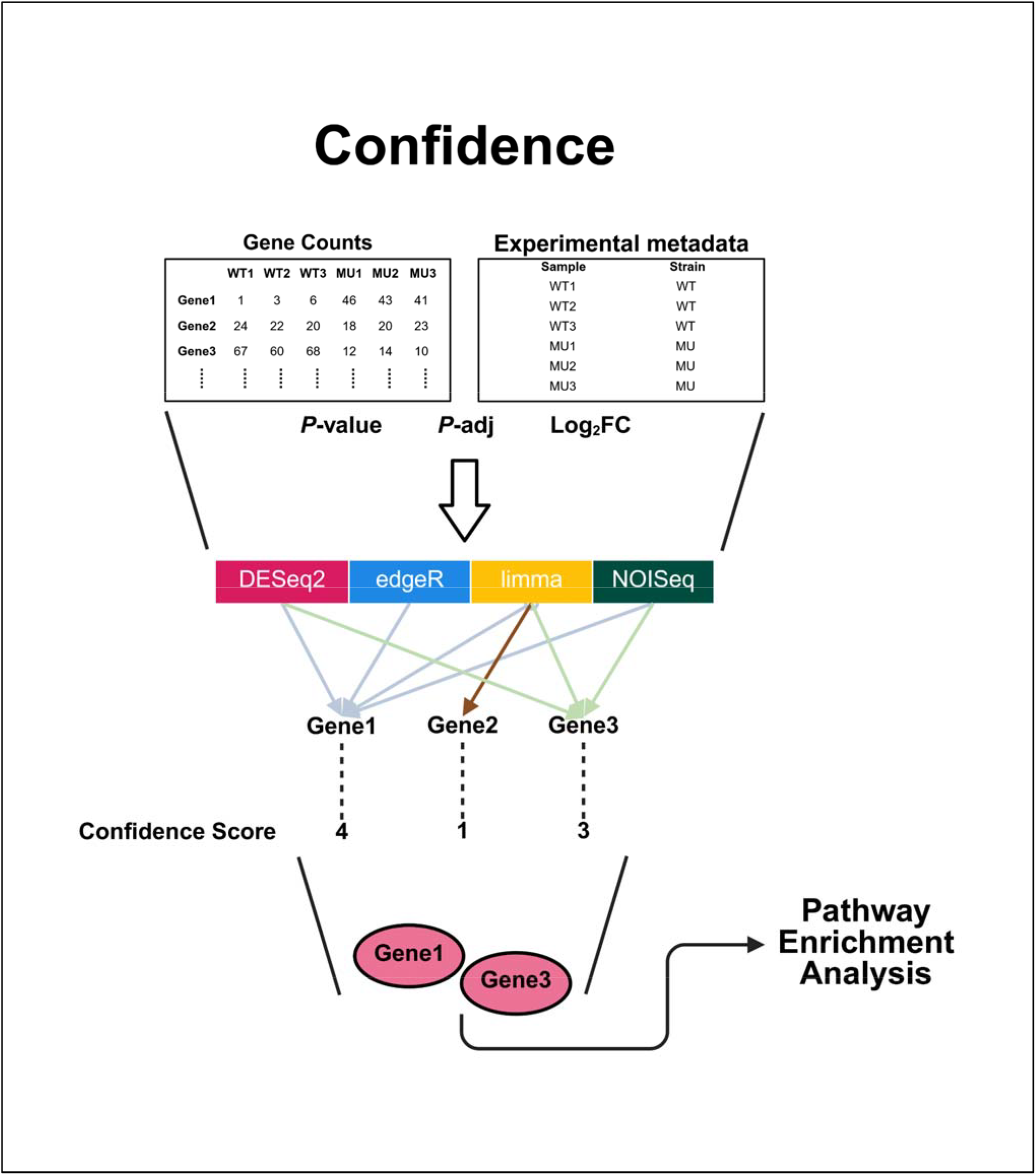

As with microarrays, differential expression analysis in RNA-seq studies compares gene counts between experimental samples through various statistical tests that were adopted from older experimental pipelines, which proved to be insufficient for this new data. In addition, variances in inter-sample library size, sequencing variability, and batch effects between biological replicates were often left unaccounted for in simple statistical analyses *(29)*. To address these issues, numerous bespoke analytical packages were developed that incorporated varying statistical assumptions about read count distributions, data normalization, and methods for hypothesis testing between samples *(30)*. In isolation, these analytical packages provide appropriate differential expression analysis based on experimental metadata and gene count input by the user. These packages can all be implemented using the R statistical programming language.

### Challenges in Differential Expression Analysis

Broadly, there are three key challenges to the use of analytical packages for differential expression analysis:

1. ***Modelling variability:*** Transcriptomic studies using deep RNA-seq often incorporate multiple biological replicates to power the experiment. Therefore, there may be variability associated with sample replicates (e.g., heterogenous disease expression between replicates), library sizes, and read quality, among other factors. Additionally, the presence of genes with abnormally skewed count distributions may cause true Differentially Expressed Gene (DEG) to be misclassified or overlooked. Since analytical packages incorporate data preprocessing measures such as batch correction, normalization, multiple test correction, and transformation to account for these factors *(31)*; because the statistical assumptions and hypothesis testing procedures differ drastically between analytical packages, the number of genes identified as differentially expressed differs between analysis packages, as does the expression level (e.g., log_2_FC, log-ratio) and significance value reported for each gene (e.g., *P*-value, *P*-adj) *(30, 32, 33)*.
2. ***Expert analysis:*** Appropriate use of analytical packages for transcriptomics requires a substantial bioinformatics and programming background. Indeed, qualitative studies have demonstrated that the complexity of software, lack of formal training, and decreased perceived ease-of-use of bioinformatics tools reduce non-bioinformaticians’ ability to perform bioinformatics analyses such as functional enrichment and/or pathways analysis *(34-36)*.
3. ***Target prioritization:*** A primary goal of transcriptomic analyses is the identification of genes whose differential regulation contributes to the experimental condition under investigation. However, the list of DEGs can be substantial. Genes of interest from these lists have traditionally been chosen arbitrarily through *P*-value, *P*-adj, and/or expression level parameters; alternatively, genes are often selected through the familiarity of the gene or its related protein to the investigator. Without a systematic, unbiased method to rank and delineate genes of interest, genes or gene sets that are reflective of real biological changes and play novel roles in the disease pathology, treatment intervention, or biological model may be overlooked.

As transcriptomics analyses incorporating next-generation RNA-seq technology have become increasingly prominent in almost every domain of biological study, it is important to provide non-bioinformaticians an avenue in which to analyze large gene expression datasets and best use the insights from these experiments. Here, we built an intuitive web-based application using Shiny, called Confidence. Confidence is easy to use and answers each of the three challenges laid out above; 1. Confidence provides a way for users to model experimental metadata and visualize the clustering of samples used in RNA-seq experiments; 2. Confidence has a simple interface that allows count.csv files, and a metadata table to be uploaded; 3. Confidence provides sequential analysis of RNA-seq derived gene counts through multiple analytical packages, providing publication-ready figures and tables that can be directly downloaded to a local machine.

## Methods

### Application Development

We built the Confidence web application using the **Shiny** package v1.10.0 *(37)* for R v4.4.3 (http://www.r-project.org), bslib v0.9.0, bsicons v0.1.2, **DESeq2** v1.44.0 *(38)*, **NOISeq** v2.48.0 *(39, 40)*, **limma** v3.60.6 *(41)*, **edgeR** v4.2.2 *(42)*, shinycssloaders v1.1.0 shinyjs v2.1.0, **plotly** v4.10.4 *(43)*, thematic v0.1.6, **ggvenn** v0.1.10 *(44)*, dplyr v1.1.4, **enrichplot** v1.24.4 *(45)*, shinyWidgets v0.9.0, spsComps v0.3.3.0, **gprofiler2** v0.2.3 *(46)*, tidyr v1.3.1, eply v0.1.2, DT v0.33, ggrepel v0.9.6, **ggplot2** v3.5.2 *(47)*, Matrix v1.7-3, SummarizedExperiment v1.34.0, Biobase v2.64.0, MatrixGenerics v1.16.0, matrixStats v1.5.0, GenomicRanges v1.56.2, GenomeInfoDb v1.40.1, IRanges v2.38.1, S4Vectors v0.42.1, BiocGenerics v0.50.0.

### Data Input

Confidence requires users to input two files: one containing gene counts and the other containing experimental metadata. Gene counts must be formatted from count estimation computational tools in a tabular format, with column values representing sample names and rows representing genes; experimental metadata can be constructed as a comma-separated values file in programs such as Microsoft Excel^®^.

### Data Modelling

Confidence provides extensive options to model experiment-associated variables found in the metadata, including selection of the main effect (the primary factor being investigated; e.g., treatment), reference class (the reference level for comparison within the main effect; e.g., untreated), and comparison class (the level to be compared to the reference class within the main effect; e.g., penicillin-treated). This is typically sufficient for two-factor comparisons (e.g., untreated vs. penicillin-treated). As a feature to guide the design of experimental questions, Confidence also allows users to build complex experimental designs controlling for multiple continuous or discrete factors (e.g., sex, age, diet) to isolate the impact of the primary variable of interest on gene expression while controlling for other factors.

### Analytical Packages

Confidence also requires the user to define a *P*-adj threshold and provides options for low count filtration based on the smallest comparison group size before initiation. The Confidence pipeline was developed to perform gene count normalization, filtration, and hypothesis testing sequentially across DESeq2 v1.44.0 *(38)*, limma v3.60.64 *(48, 49)*, edgeR v4.2.2 *(42)*, and NOISeq v2.48.0 *(39, 40)*, which are the most popular analytical packages used in published transcriptomic research. The default normalization settings and hypothesis tests were maintained in each analytical package (see **Table 1**) as per their respective vignettes *(38-42, 48)*. Annotation of Ensembl gene IDs was facilitated using the biomaRt database v2.60.1 *(50-52)*. A dropdown list of Human, Rat, and Mouse genome annotations are provided to characterize ENSEMBL gene IDs. This list will be expanded in the future to include more species. Due to the presence of uncharacterized mRNA reads in RNA-seq datasets, unannotated results that do not have gene names are also included in the Confidence results table, relying on their ENSEMBL gene ID annotation instead. An overview of the Confidence application methodology is presented in **Figure 1**.

**Table 1.**

| Platform | Original Use | Proportion of False Discoveries |  | Design Formula Complexity | Normalization; Transformation | Multiple test correction method | Market share |
| --- | --- | --- | --- | --- | --- | --- | --- |
|  |  | Small Sample Sizes | Large Sample Sizes |  |  |  |  |
| **DESeq2** | RNA-seq, CHIP-seq | Low | Low | Multi-factor | DESeq2 Size-Factors;<br>**Variance Stabilizing Transformation**, Regularized Log Transformation | Holm, Hochberg, Hommel, Bonferroni, **Benjamini-Hochberg**, Benjamini-Yekutieli | 26.36% |
| **limma** | Microarrays | Low | Low | Multi-factor | TMM;<br>voom | Holm, Hochberg, Hommel, Bonferroni, **Benjamini-Hochberg**, Benjamini-Yekutieli | 3.13% |
| **edgeR** | RNA-seq | Moderate | Moderate | Single factor | **TMM**, Upper quartile, DESeq2-like;<br>Counts per million | **Benjamini-Hochberg**, Benjamini-Yekutieli, Holm | 20.84% |
| **NOISeq** | RNA-seq | Moderate-high | Low-Moderate | Single factor | **RPKM**, TMM, Upper quartile;<br>None | a posteriori probability-akin to Benjamini-Hochberg FDR | 1.38% |

### Differential Gene Expression Analysis

After contrasting samples based on selected main effects, each analytical package outputs a table of DEGs along with each gene’s ID, log_2_FC, *P*-value, and *P*-adj, across conditions. NOISeq is an exception, however, as it produces an *a posteriori* probability *p* that a gene is differentially expressed between two conditions *(39, 40)*. Here, we specify that the *P*-adj for NOISeq is equivalent to 1-*p*, as instructed in the NOISeq vignette.

### Confidence Score

After sequential tests of differential expression are performed by each analytical package, Confidence Scores are calculated for each DEG. The Confidence Score is a tally of the number of packages where a significant DEG is demonstrated to be up-or down-regulated. This ‘consensus’ among analytical packages allows for the filtering of genes that are likely to be both biologically and statistically significant. As an example, if three of the four packages implemented in Confidence report a DEG, the gene’s associated Confidence Score is 3. Herein, we qualitatively describe genes with a Confidence Score of 0 as ‘not confident’, Confidence Score of 1 as ‘least confident’, a Confidence Score of 2 as ‘moderately confident’, a Confidence Score of 3 as ‘highly confident’, and a Confidence Score of 4 as ‘most confident’. To elucidate overlaps between DEGs reported by each analytical package, a four-way Venn diagram is automatically generated by Confidence on the named list using the ggvenn package 0.1.10 *(53)*.

### Outputs and Visualizations

Confidence allows for the generation of standard visualizations found in transcriptomic literature. Principal component analysis (PCA) was integrated into Confidence using the DESeq2 plotPCA function and formatted using ggplot2 *(55)*. PCA was performed using the *prcomp* function on variance stabilizing transformed gene counts; a PCA biplot is immediately available when experimental metadata and gene count files are submitted, so that the effect of including/excluding experimental variables and metadata factors (e.g., sex, diet, outliers, and low gene counts) can be visualized through sample clustering. To investigate the spread of DEGs according to their log_2_FC and *P*-adj values, volcano plots are generated on DEGs from each analytical package using ggplot2 v3.5.2 and plotly v4.10.4. Additionally, Box plots can be generated for each DEG to visualize normalized gene counts between experimental conditions; for box plots only, normalization is performed using log_2_-transformed, DESeq size-factors. These box plots are embedded into the final Confidence results table and are immediately accessible through a clickable button.

### Pathway Analysis

Pathway enrichment analysis (PEA) tools were additionally programmed into Confidence using the *GOSt* function from the gprofiler2 package v0.2.3 *(56)*. *GOSt* analysis is provided to perform PEA using numerous databases including gene ontology (GO), Kyoto Encyclopedia of Genes and Genomes (KEGG), Reactome (REAC), WikiPathways (WP), TRANScription FACtor database (TF), MicroRNA-Target database (miRTarBase), Human Protein Atlas (HPA), Comprehensive Resource of Mammalian protein complexes (CORUM), and Human Phenotype Ontology (HP), simultaneously.

## Results

Confidence can be found, here: https://confidence.apps.meds.queensu.ca/

To showcase Confidence, we have included sample figures that were generated in Confidence using toy data (**Figure 2**). Confidence allows for modelling of metadata (by treatment, animal ID, sample number, and sample ID; **Figure 2A-D**), an understanding of how many DEGs each package identified using a Volcano plot (**Figure 2E**) and how many intersected (**Figure 2F**), and the expression profiles of different, user defined genes (**Figure 2G/H**). Confidence also visualizes the functional analysis of the gene list as an interactive dot plot of all enriched terms organized by domain (**Figure 3A**), as well as user-defined dot plots of the top functions/processes/targets in each domain (**Figure 3B**).

**Figure 2.**
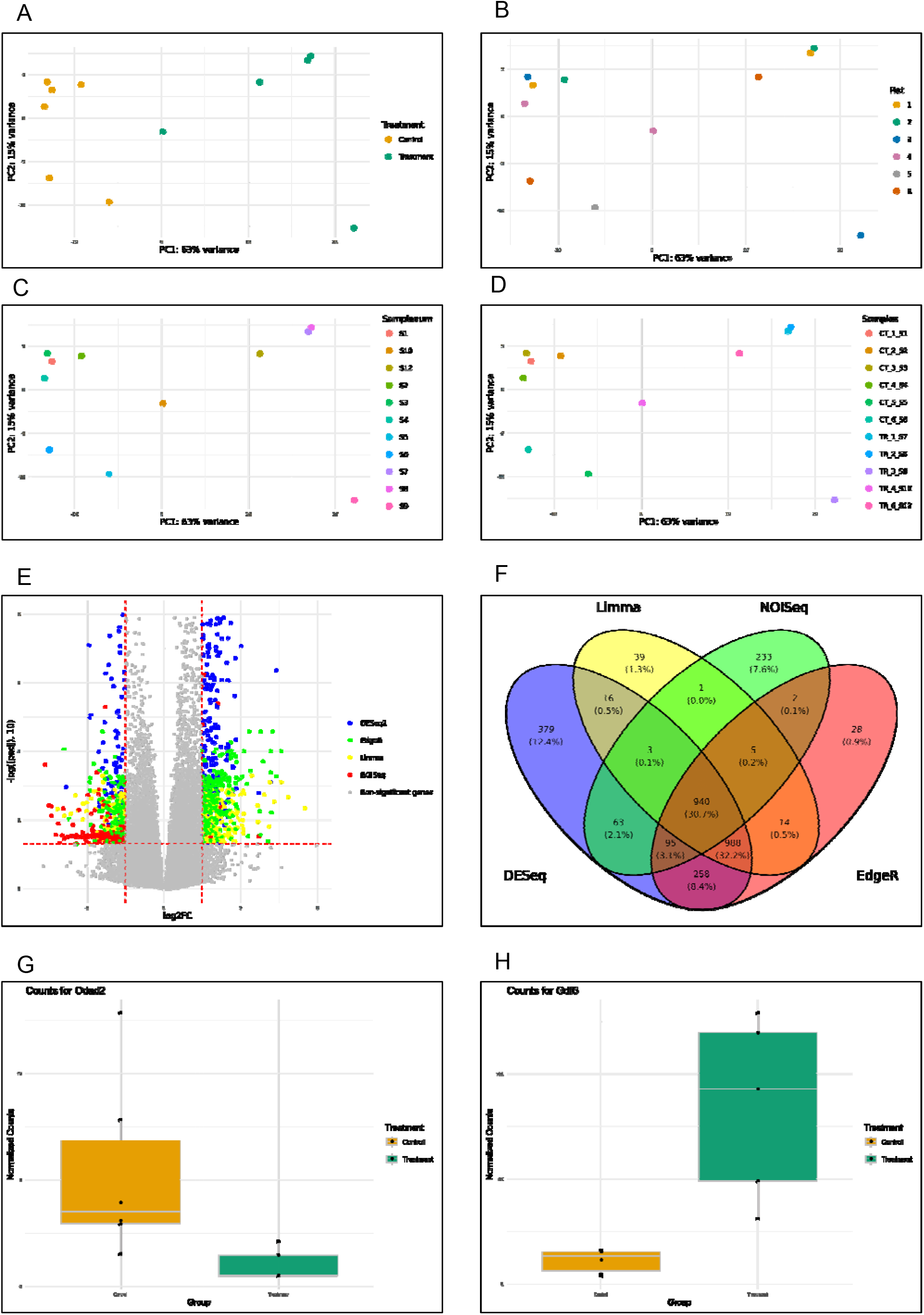

**Figure 3.**
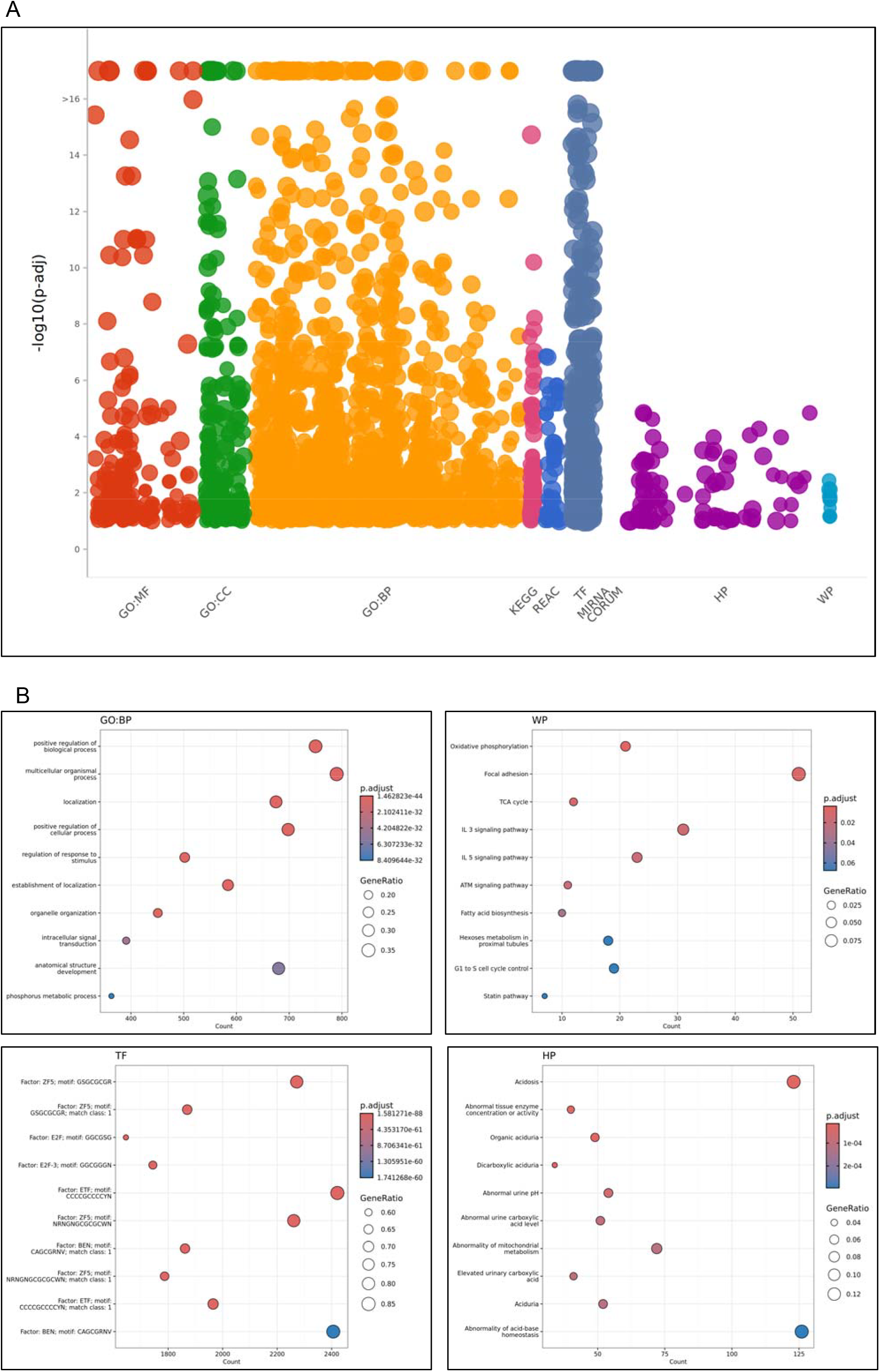

## Discussion

Differential expression analysis is often the penultimate step of transcriptomic studies, which statistically evaluates the differences in gene counts between experimental groups. Since the publication of the first RNA-seq study in 2006 *(57)*, several analytical packages and methodologies have been developed to normalize, model, and test differential expression effectively *(58)*. However, large variations in the number of discovered genes, false discovery rate, and expression changes across these analytical packages may confound the interpretation of results *(30, 32, 33, 59-62)*. Among these transcriptomics analytical packages, DESeq2, limma, edgeR, and NOISeq are commonly used in deep RNA-seq studies *(62-64)*. Additionally, DESeq2 accounted for 26.36%, edgeR for 20.84%, limma-voom for 3.13%, and NOISeq for 1.38% of all citations for RNA-seq studies *(58)*. NOISeq, the most popular non-parametric method, was implemented into Confidence as it offers a unique computational and theoretical approach relative to parametric methods such as DESeq2, edgeR, and limma.

Here, we introduced the Confidence application, which embeds each of these analytical packages into a single analysis pipeline to leverage their unique features in a user-friendly application. The use of multiple analytical packages to simultaneously assess gene expression has previously been implemented in other tools, including *consensusDE* and *consexpression (65, 66)*; however, these tools both require the use of programming languages.

In untargeted omics analyses more broadly, the process to select features to characterize, validate, or manipulate must take several considerations into account, including the cost of reagents and the considerable amount of time needed to investigate findings. We propose that Confidence can provide an intuitive means to filter large gene lists and streamline gene selection. Additionally, although many analytical package-consensus tools have been developed *(65, 66)*, they are implemented as computational packages within coding languages such as Python or R. In a survey conducted by Alomair *et al*. (2023) of scientists, only one-third used programming languages to address biological problems, and half reported hiring a bioinformatician to conduct the analysis *(34)*. To address this issue, Confidence was developed as a web-based application, which included options to produce the visualizations presented in this study, as well as clear descriptions of analytical methods, plot interpretations, and plot customization tools.

### Confidence addresses three major challenges to identify and prioritize DEGs from transcriptomic studies

1. Confidence allows the easy modelling of data using different sample attributes reported in the user-defined metadata; PCA’s will allow for the rapid review of sample variability using condition, as well as other information such as sex, batch, animal ID, and time.
2. Confidence integrates standard pipelines for differential expression analysis into a simple web-based platform, which may ensure that differential expression analysis is more accessible for investigators without programming experience.
3. Using the Confidence Score, Confidence provides a consensus tool to identify common DEGs across analytical packages employing diverse statistical methodologies, enhancing target prioritization.

## Conclusions

We have developed the Confidence application to perform multi-package differential expression analysis, provide intuitive scoring systems to streamline gene prioritization and selection, and provide a web-based tool for non-bioinformaticians to analyze transcriptomics data. Further developing Confidence to integrate the vast array of analytical packages available in other omics fields, such as proteomics and metabolomics, and to gauge their performance, can help identify novel markers implicated in disease processes. Finally, we foresee that the ability for Confidence to streamline marker selection without the influence of knowledge bias by the investigator will reduce costs associated with unproductive validation and experimentation in transcriptomics and beyond.

## Acknowledgments

We would like to thank the MitoMetabLab at Queen’s University for their suggestions and tests of the Confidence application. We would also like to thank the Queen’s CardioPulmonary Unit (QCPU), and the Translational Institute of Medicine (TIME) at Queen’s University.

## Author contributions

Conceptualization: A.S., B.P.O, K.DS., AP., C.C.T.H.

Methodology: A.S., B.P.O., K.DS., C.C.T.H.

Investigation: A.S.

Visualization: B.P.O., A.S.

Formal Analysis: A.S., B.P.O

Funding acquisition: K.DS., C.C.T.H.

Project administration: A.S., B.P.O., K.DS., C.C.T.H.

Resources: B.P.O., M.S., K.DS., C.C.T.H.

Software: B.P.O., A.S., C.C.T.H.

Supervision: K.DS., C.C.T.H.

Writing – original draft: A.S., K.DS., C.C.T.H.

Writing – review & editing: A.S., B.P.O, K.DS., C.C.T.H.

## Data and materials availability

The Confidence application is available at https://confidence.apps.meds.queensu.ca/. Any questions regarding code availability and the conclusions of this paper should be directed to the corresponding author.

